# Distinct isoforms of Nrf1 diversely regulate different subsets of its cognate target genes

**DOI:** 10.1101/356071

**Authors:** Meng Wang, Lu Qiu, Xufang Ru, Yijiang Song, Yiguo Zhang

**Author notes:** Correspondence should be addressed to Yiguo Zhang or).

## Abstract

The single *Nrf1* gene has capability to be differentially transcripted alongside with alternative mRNA-splicing and subsequent translation through different initiation signals so as to yield distinct lengths of polypeptide isoforms. Amongst them, three of the most representatives are Nrf1α, Nrf1β and Nrf1γ, but the putative specific contribution of each isoform to regulating ARE-driven target genes remains unknown. To address this, we have here established three cell lines on the base of the Flp-In^™^ T-REx^™^ system, which are allowed for tetracycline-inducibly stable expression of Nrf1α, Nrf1β and Nrf1γ. The RNA-Sequencing results have demonstrated that a vast majority of differentially expressed genes (i.e. >90% DEGs detected) were dominantly up-regulated by Nrf1α and/or Nrf1β following induction by tetracycline. By contrast, other DEGs regulated by Nrf1γ were far less than those regulated by Nrf1α/β (i.e. ~11% of Nrf1α and 7% of Nrf1β). Further transcriptomic analysis revealed that tetracycline-induced expression of Nrf1γ significantly increased the percentage of down-regulated genes in total DEGs. These statistical data were further validated by quantitative real-time PCR. The experimental results indicate that distinct Nrf1 isoforms make diverse and even opposing contributions to regulating different subsets of target genes, such as those encoding 26S proteasomal subunits and others involved in various biological processes and functions. Collectively, Nrf1γ acts as a major dominant-negative competitor against Nrf1α/β activity, such that a number of DEGs regulated by Nrf1α/β are counteracted by Nrf1γ.

## Introduction

Nuclear factor-erythroid 2-related factor 1 (Nrf1) acts as a transcription factor belonging to the cap’n’collar (CNC) basic-region leucine zipper (bZIP) family, which is indispensable for maintaining cellular homoeostasis and organ integrity during normal development and growth, as well as the adaptation to other pathophysiological processes [1–3]. It is important to note that the unique function of Nrf1 is finely tuned by a steady-state balance between production of the CNC-bZIP protein (i.e. translation of transcripts) and its concomitantly (i.e. post-transcriptional and post-translational) processing to give rise to distinct multiple isoforms (called proteoforms, with different and even opposing abilities) before being turned over. These distinct proteoforms of Nrf1 are postulated to together confer on the host robust cytoprotection against a vast variety of cellular stress through coordinately regulating distinctive subsets of important homoeostatic and developmental genes. The transcriptional expression of such key genes are driven by antioxidant response elements (AREs) and/or other *cis*-regulating consensus sequences, some of which are conversed with the activating protein-1 (AP-1) binding site, within these gene promoter regions.

The single gene of *Nrf1* (also called *Nfe2l1*) is allowed for differential transcriptional expression to yield multiple mRNA transcripts (between ~1.5 kb and ~5.8 kb) and subsequently alternative translation into distinct polypeptide isoforms (between ~25 kDa and ~140 kDa), which are determined to be differentially distributed in embryonic, fetal and adult tissues, including liver, brain, kidney, lung, heart, skeletal muscle, bone, testis, ovary, placenta and others [4–7]. Amongst such isoforms, the full-length Nrf1 (designated Nrf1α) is yielded by the first translation initiation signal within the main open reading frame of alternatively spliced mRNA transcripts, in which the exon 4 [encoding the peptide ^242^VPSGEDQTALSLEECLRLLEATCPFGENAE^271^, that was named the Neh4L (Nrf2-ECH homology 4-like) region] is removed from its long isoform TCF11 (transcription factor 11) in the human [5]. Albeit Nrf1 lacks the Neh4L region, it was identified to retain a strong transactivation activity largely similar to TCF11 [8].

By contrast with Nrf1α, the short isoform Nrf1β [which was early designated as LCR-F1 (locus control region-factor 1)] is determined to be generated through the in-frame translation that is initiated by an internal perfect Kozak’ starting signal (5’-puCCATGG-3’), which is situated within and around the four methionine codons between positions 289 and 297 in the mouse [4, 5, 9]. By bioinformatic analysis, it is thus inferred that Nrf1β lacks the N-terminal domain (NTD, aa 1-124) and its adjacent acidic domain 1 (AD1, aa 125-296) [10, 11]. Later, Nrf1β is also determined to exhibit a weak transactivation activity [6, 12–14], but stimulation of Nrf1β activity appears to be dependent on distinct stressors that had been administrated in different cell lines [13–15]. Furthermore, a small dominant-negative isoform, called Nrf1γ [12,13], is produced by the potential in-frame translation starting at the putative methionine of position 584, as well as by the putative endoproteolytic processing of longer Nrf1 proteins. When generation of Nrf1γ is blocked, the transactivation activity of Nrf1β is significantly increased [12]. On the contrary, when Nrf1γ is forcedly expressed, the consequence enables for a possible interference with the functional assembly of each of the active transcription factors (i.e. Nrf1α or Nrf2) with its heterodimmeric partner (i.e. sMaf and other bZIP proteins), in order to down-regulate expression of AP1-like ARE-driven target genes [12, 13].

To date, it is, however, unknown how each isoform of Nrf1 contributes to its unique role in regulating expression of ARE-driven cytoprotective genes against various physiopathological stresses. To address this issue, we have here created distinct three tetracycline (Tet)-inducibly stable expression cell lines, each of which exhibits Nrf1α, Nrf1β, and Nrf1γ, respectively, with similarities and differences of structural domains as shown diagrammatically (Fig. 1A). Subsequently, different changes in the transcriptomic expression mediated by Nrf1α, Nrf1β, or Nrf1γ were analyzed by RNA-Sequencing (RNA-Seq), some of which were further validated by quantitative PCR experiments. Collectively, we first discovered that both Nrf1α and Nrf1β make main contributions to Nrf1-mediated transcriptional expression of cognate target genes. A vast majority of differentially expressed genes (DEGs) are up-regulated by Nrf1α and/or Nrf1β upon either stable expression induced by Tet treatment. By sharp contrast, an array of similar or different DEGs regulated by Nrf1γ are far less than those genes regulated by Nrf1α or Nrf1β. This demonstrates that Nrf1γ exerts a distinguishable effect from the other two isoforms Nrf1α/β, on the transcriptional expression of Nrf1-target genes, because its inducible expression by Tet significantly increased the percentage of down-regulated genes among DEGs detected. Collectively, this work provides a further understanding of distinctions in the transcriptional regulation of its different isoforms towards Nrf1-target genes.

**Figure 1.**
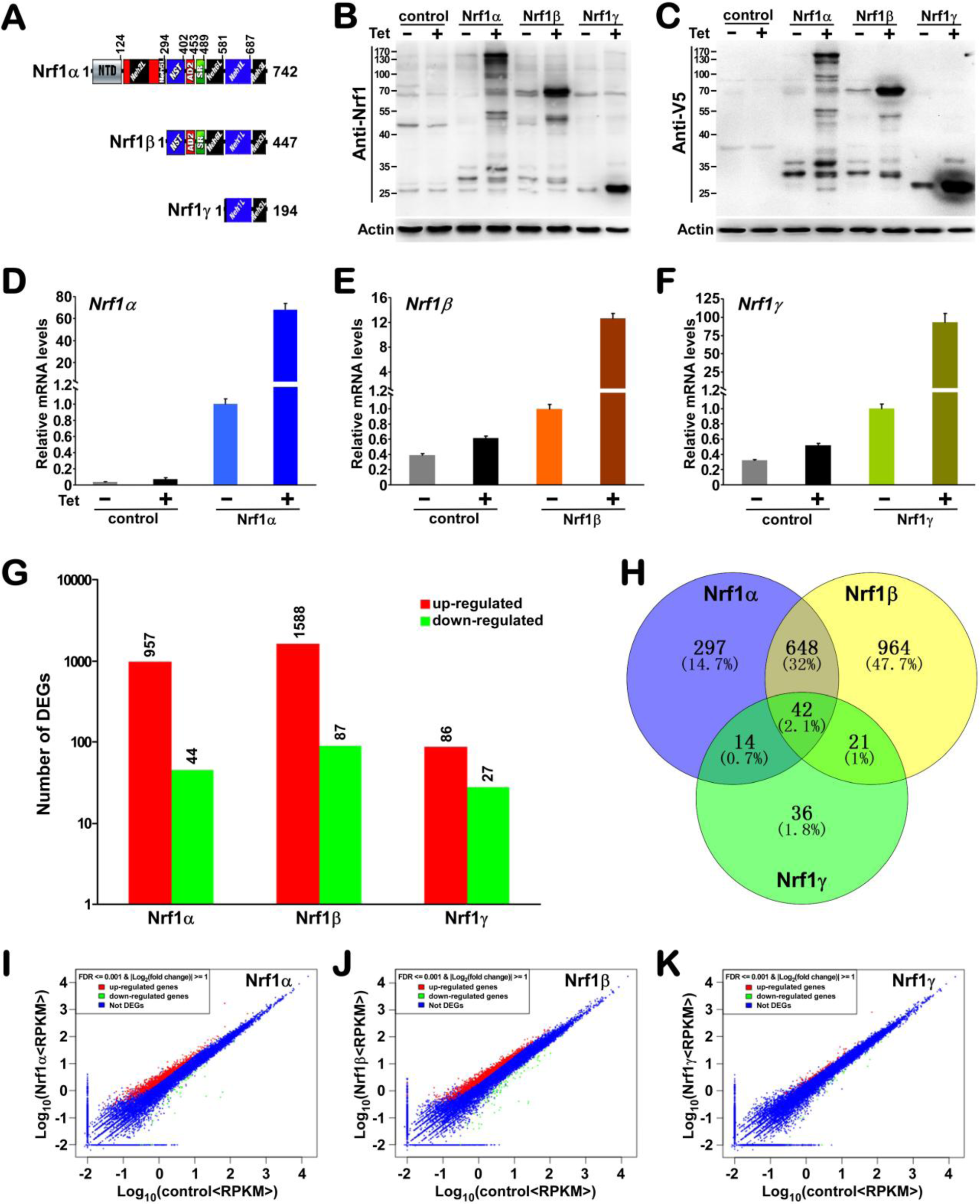
Identification of distinct Nrf1 isoforms-expressing cell lines along with global statistical analysis of RNA-Seq data. (A) Shows schematics of the similarities and differences between structural domains of Nrf1α, Nrf1β, and Nrf1γ. (B, C) Total lysates of each cell lines that had been treated with 1 μg/ ml of tetracycline (+) or not (-) were subjected to protein separation by 12% Laemmli SDS-PAGE gels running in the pH 8.9 Tris-glycine buffer, and visualization by immunoblotting with Nrf1 or V5 antibody to identify Tet-inducible expression of distinct Nrf1 isoforms. (D to F) These cell lines were further identified by quantitative real-time PCR. (G) Shows an overview of differentially expressed genes (DEGs) indicated in the stat chart of RNA-seq data. (H) Shows the Venn chart with the common or unique DEGs among sequenced samples after being normalized by control sample. (I to K) Pairwise scatter plots are used to identify global changes and trends in gene expression among three conditions.

## Results

### Identification of differentially expressed genes in distinct isoforms-expressing cell lines by RNA-Sequencing

To gain an in-depth insight into distinct contributions of Nrf1 isoforms (i.e. Nrf1α, Nrf1β and Nrf1γ) to the precision regulation of different subsets of target genes, each of these isoform-expressing cell lines was herein established by Flp recombinase-mediated integration on the base of the Flp-In^TM^ T-REx^TM^-293 host cells. These cell lines had been transfected with each of pcDNA5/FRT/TO-V5 expression constructs for distinct cDNA sequences encoding Nrf1α, Nrf1β and Nrf1γ (as shown schematically in Fig. 1A) before being selected, whilst the empty expression vector-transfected host cells served as a negative control in the parallel experiments. The inducible expression of Nrf1α, Nrf1β and Nrf1γ was under control by tetracycline (Tet) for distinct experimental requirements, before which the positively-selected clones of cell lines were maintained within double antibiotics 150 μg/ml hygromycin B and 15 Mg/ml blasticidin. Of note, the stable expression of distinct isoforms was monitored by interaction of Tet with its repressor TetR leading to release from the Tet operator and then induction of interested gene transcription.

As anticipated, after 12-h treatment of HEK293C^Nrf1α^, HEK293D^Nrf1β^ and HEK293E^Nrf1v^ cell lines with 1 μg/ml of tetracycline, they were harvested, followed by immunoblotting (Fig. 1, B & C) and quantitative PCR (Fig. 1, D to F) to determine their protein and mRNA levels, respectively. The immunoblotting with antibodies against Nrf1 and its C-terminal V5 tag showed that the inducible expression of Nrf1α, Nrf1β and Nrf1γ were validated by treatment of tetracycline in their respective cell lines. Nrf1α was exhibited as two major close isoforms of ~140/130-kDa (i.e. glycoprotein and deglycoprotein) alongside with two minor processed isoforms of ~100/90-kDa (Fig. 1B, C), Nrf1β displayed as a major ~70-kDa protein and another minor processed ~50-kDa polypeptide (called Nrf1β2), whilst Nrf1γ expressed as a minor ~36-kDa protein and a major processed ~25-kDa polypeptide (also called Nrf18).

These Tet-inducibly isoform-expressed and negative control cell lines were subjected to extraction of total RNAs before being sequenced (i.e. RNA-Seq). An overview of the primary sequencing data was then depicted in Table S1. Following the processing and analysis of the RNA-Seq data, the gene expression levels were calculated by the RPKM (*Reads Per kb per Million reads*) method [16]. Of note, the DEGs were screened out, with the threshold of *P*-value in multiple tests at a false discovery rate (FDR) ≤ 0.001 [17], along with an absolute value of Log_2_ (fold change) ≥ 1, and identified by calculating each gene expression in sample groups versus controls as indicated in the Stat Chart (Fig. 1G). By comparison to control cells, 1001 genes showed significant changes (Table S2) in the transcriptional expression as accompanied by stable Nrf1α-expression in Tet-inducible HEK293C^Nrf1α^ cells, of which 957 genes (i.e. 95.6%) were up-regulated (Fig. 1G, *red bar*) and other 44 genes (i.e. 4.4%) were down-regulated (*green bar).* And, transcriptional expression of as many as 1675 genes was also significantly regulated upon stable expression of Nrf1β in the Tet-inducible HEK293D^Nrf1β^ cells, 94.8% of which (i.e. 1588 genes) were up-regulated and other 5.2% (i.e. 87 genes) were down-regulated (Fig. 1G, and also Table S3). By striking contrast, only 113 genes were significantly transcriptionally changed in the stable Tet-inducibly Nrf1γ-expressing HEK293E^Nrf1γ^ cells. This appeared to be just about one-tenth of the number of those genes measured in either HEK293C^Nrf1α^ or HEK293D^Nrf1β^ cells. Amongst the 113 genes regulated by Nrf1γ, besides only 27 genes (i.e. 23.9%) were down-regulated, the other 86 genes (i.e. 76.1%) were still up-regulated, (Fig. 1G, and Table S4). This implies a possible counteraction of Nrf1γ competitively against Nrf1α/β down-regulation of some genes.

After normalization by the control, each isoform-specific or their common DEGs amongst three Nrf1 isoforms were statistically shown with the Venn diagram (Fig. 1H). Their scatterplots were useful to identify global changes with distinct trends in gene expression between pairs of conditions (Fig. 1, I to K). Overall, these data demonstrate that Nrf1β regulates the greatest number of DEGs amongst these three isoforms, followed by Nrf1α, whereas the number of DEGs regulated by Nrf1γ is far less than that of DEGs regulated by either of the former two isoforms.

### Functional annotation of differentially expressed genes regulated by each of distinct Nrf1 isoforms

Further data analysis of thousands of DEGs involved in distinct biological processes is an important downstream task following RNA-Seq to understand relevant meanings of those ‘interested’ genes regulated by each isoform of Nrf1. Of note, gene ontology (GO) is an internationally-standardized functional classification system of genes, which offers a dynamically-updated controllable vocabulary with a strictly defined concept to describe comprehensively properties of distinct genes and their products in any organism, and covers three major domains, including cellular component, molecular function and biological process. The enrichment analysis of GO provides all relevant terms that are significantly enriched in the DEGs, by comparison to the genomic background, and then filters the DEGs that correspond to potential biological functions. The *d*atabase for *a*nnotation, *v*isualization and *i*ntegrated *d*iscovery (DAVID) of bioinformatic resources consists of an integrated biological knowledgebase with analytic tools, which is used for systematic extraction of biological features and/or meanings associated with large lists of genes [18]. Therefore, in order to investigate the relationship and difference in gene regulation by amongst these three Nrf1 isoforms, the enriched functional annotation of GO terms (Tables S5 to S7) was mapped with the DEGs regulated by each isoform. The involved DEGs in different terms were identified by the DAVID tool and listed according to their enrichment P-values. Those genes are then enabled for an interaction with each other to play some key roles in the certain biological functions. Thus, the results of pathway enrichment analysis (Tables S8 to S10) was obtained on the base of the major public pathway-related database KEGG (Kyoto Encyclopedia of Genes and Genomes [19]). This is helpful to further understand biological functions of distinct subsets of genes regulated by each Nrf1 isoform.

The top 20 of significantly enriched functional annotation terms (Fig. 2) are represented by different GO classes and KEGG pathways. Within the GO-classified biological processes, the Nrf1α-regulated DEGs are highly enriched in terms of transcriptional regulation, RNA metabolism, cell cycle, macromolecule catabolic process, cellular response to stress, protein localization, metabolic process and chromosome organization (Fig. 2A). Almost all these enriched biological process terms of Nrf1α-regulated DEGs were also significantly enriched in other DEGs regulated by Nrf1β. Besides similar terms, Nrf1β-regulated DEGs were also involved in intracellular signaling cascade, phosphorylation, transcriptional regulation of RNA polymerase-II promoter and biosynthetic process (Fig. 2E). By contrast, just the fewer numbers of DEGs were regulated by Nrf1γ, and hence only a few of genes were mapped to distinct groups of biological processes, such as regulation of cell proliferation, response to wounding, protein metabolic process, blood vessel development, peptide cross-linking and plasma membrane long-chain fatty acid transport (Fig. 2I). As for cellular components, those DEGs regulated by Nrf1α and Nrf1β are also highly enriched in terms of non-membrane -bounded organelles, membrane-enclosed lumen, organellic lumen, juxtanuclear lumen, nucleoplasm, nucleolus, cytosol, cytoskeletal part, Golgi apparatus, endomembrane system and extrinsic to membrane (Fig. 2, B & F). These DEGs were also highly enriched for ion-binding, nucleotide-binding, ATP-binding and transcription regulator activity in molecular function groups (Fig. 2, C & G). Notably, there are more about 30 % of DEGs regulated by Nrf1β (versus control) mapped to each of the same GO term as that regulated by Nrf1α, implying that both regulate distinct subsets of more Nrf1-target genes. By sharp contrast, only a few of genes regulated by Nrf1γ were mapped to couple cellular component groups, like non-membrane-bounded, organelle, membrane-enclosed lumen and nucleolus (Fig. 2J). Meanwhile, the Nrf1γ-regulated DEGs were also not significantly enriched in molecular function terms (Fig. 2K). As such, not all of the GO terms mapped with Nrf1γ-regulated DEGs also appear in the former two groups (see tables S5 to S7), implying that several genes regulated by Nrf1γ are distinguishable from those regulated by Nrf1α/β.

**Figure 2.**
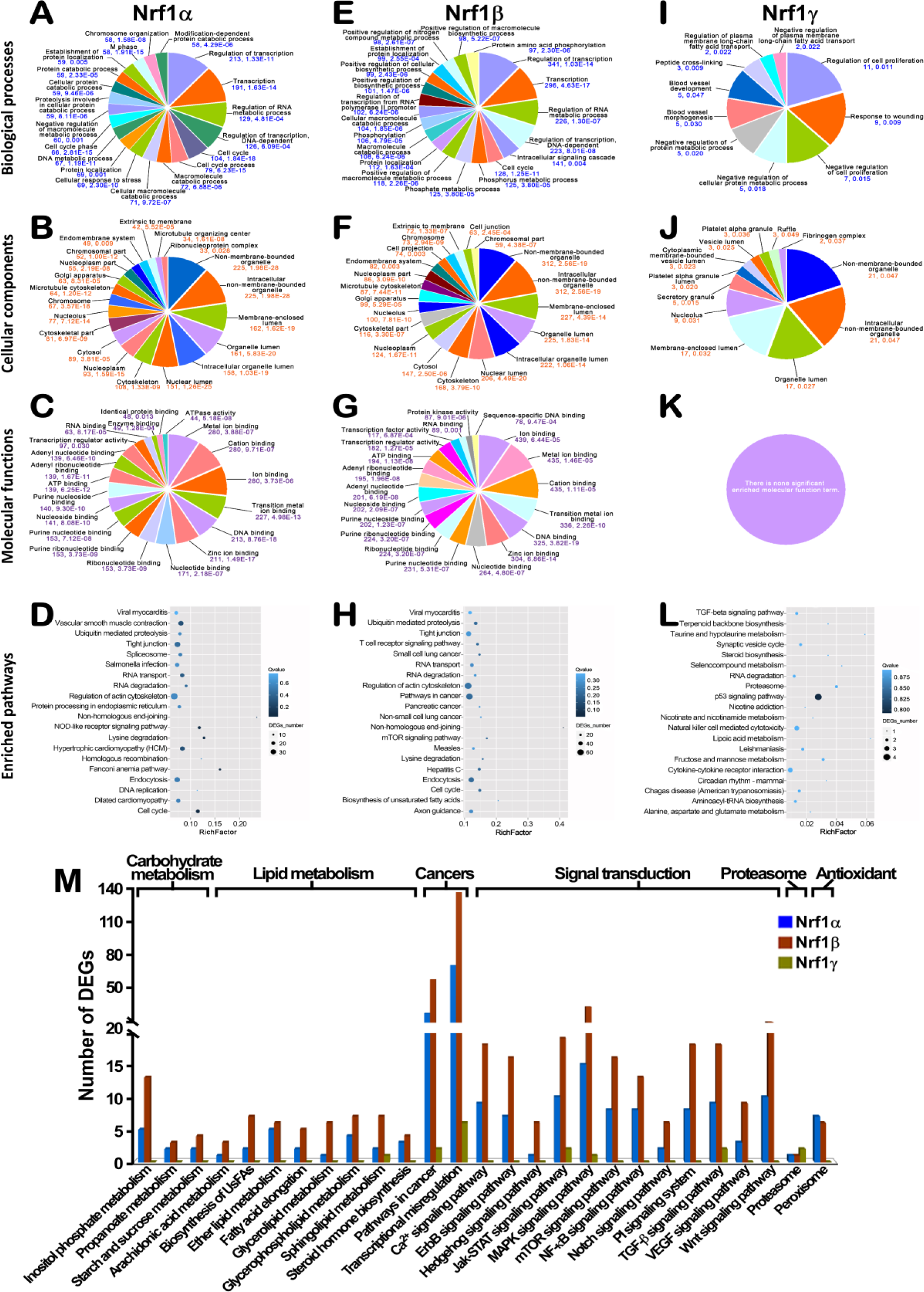
Functional annotation of DEGs regulated by three Nrf1 isoforms. By GO analysis of Nrf1α-regulated DEGs, multilevel distribution is shown for the GO categories: “biological process” (A), “cellular component” (B) and “molecular function” (C). Similar categories were shown in the cases of Nrf1β-regulated DEGs (E to G) and Nrf1γ-regulated DEGs (I to K). The significant enriched GO terms presented in the pie charts are ranked by the number of DEGs, and the numerical values below each term are represented by the number of DEGs and P-value. The top-ranked 20 pathways enriched according to the *Q* value were plotted for Nrf1α-regulated DEGs (D), Nrf1ß-regulated DEGs (H), and Nrf1γ-regulated DEGs (L). In the scatter plots, the rich factor is a ratio of DEGs’ number annotated in one pathway term to all genes’ number annotated in this pathway term. The *Q* value is corrected P-value ranging from 0 to 1, and less *Q* value means greater intensiveness. Furthermore, the number of DEGs mapped to the pathways, that were associated with carbohydrate metabolism, lipid metabolism, cancers, signal transduction, proteasome and redox signaling, was calculated in the stat chart (M).

Furtherly, the results obtained from relevant pathway enrichment analysis (Tables S8 to S10) revealed that DEGs regulated by three Nrf1 isoforms are highly enriched in the terms of human diseases (i.e. dilated cardiomyopathy, hepatitis C, hypertrophic cardiomyopathy, measles, non-small cell lung cancer, pathways in cancer, pancreatic cancer, salmonella infection, small cell lung cancer, viral myocarditis), metabolism (e.g. alanine, aspartate and glutamate metabolism, biosynthesis of unsaturated fatty acids, inositol phosphate metabolism, lysine degradation, purine metabolism) and genetic information processing (e.g. DNA replication, homologous and non-homologous end-joining recombination, protein processing in the endoplasmic reticulum, ubiquitin-mediated proteolysis, RNA transport and degradation, and spliceosome). These DEGs are also involved in pathways of cellular processes (e.g. cell cycle, endocytosis, regulation of actin cytoskeleton and tight junction), organismal systems (e.g. axon guidance, NOD-like receptor signaling, T cell receptor signaling, vascular smooth muscle contraction) and environmental information processing (e.g. mTOR signaling, and TGF-β signaling). Collectively, Figure 2(D, H & L) showed the top 20 of statistical pathway enrichment within DEGs regulated by each isoform of Nrf1. Of note, Nrf1 is also essential for maintaining cellular homoeostasis and organ integrity in multifunctional responses to a variety of endogenous and exogenous stimulators during normal development and growth. Thus, the number of DEGs was further calculated and thus mapped to distinct pathways responsible for carbohydrate metabolism, lipid metabolism, cancers, signal transduction, proteasome and redox signaling (Fig. 2M). These results demonstrate distinct Nrf1 isoform-specific regulation of different subsets of target genes, which are diversely involved in multiple physio-pathological processes. Further functional annotation indicates that three Nrf1 isoforms have many similarities in exerting certain biological functions, but there still exist a few of differences amongst these three isoforms. They are required for coordinated cooperation together in a given organism to play more similarly overlapping, but also different and even opposing, biological functions of Nrf1. For example, Nrf1α and Nrf1β, but not Nrf1γ, may make main contributions to the physiological function of Nrf1, whilst Nrf1γ acts, at least in part, as a dominant-negative inhibitor of Nrf1.

### Distinct expression profiles of the top DEGs regulated by Nrf1α, Nrf1β and Nrf1γ

To investigate distinctions in the target gene expression patterns regulated by between Nrf1α, Nrf1β and Nrf1γ, the differentially expressed data of each isoform-specific and/or their sharing top 10 of RPKM DEGs were extracted (Table S11). These unique and/or common DEGs were shown in the heatmap (Fig. 3A) and also clustered by HemI [20] (Fig. 3B). According to both RPKM and fold changes in gene expression, the unique and/or some common top-ranked DEGs were picked out to build a gene-regulatory network among three Nrf1 isoforms (Fig. 3C, and Table S12). As shown in the heatmap, all the top 10 RPKM Nrf1α-specific genes showed up-regulation, whilst 9 of the top 10 RPKM Nrf1β-specific genes were up-regulated, but with only one gene (i.e. KRT18) being down-regulated by Nrf1β. By contrast, 6 of the top 10 RPKM Nrf1γ-specific genes were up-regulated, whereas other 4 genes were down-regulated by this isoform (Fig. 3A). The top 10 RPKM Nrf1α/β-shared regulatory genes were up-regulated by both Nrf1α and Nrf1β, but not by Nrf1γ, whilst the top 10 RPKM Nrf1α/γ-coordinated genes were up-and down-regulated at a 5:5 ratios by Nrf1α and Nrf1γ rather than Nrf1β (Fig. 3A). In addition, 9 of the top 10 RPKM Nrf1β/γ-shared regulatory genes showed up-regulation, but only one (i.e. *TNFRSF12A*) was down-regulated. Another 9 of the top 10 RPKM genes were commonly up-regulated by three Nrf1 isoforms, with only an exception of *LCP1* (lymphocyte cytosolic protein 1) that was significantly down-regulated by Nrf1β but up-regulated by other two isoforms (Fig. 3A). Collectively, the global statistical results of DEGs (as described above in Fig. 1F) also revealed that Nrf1β specifically regulated a maximum number of DEGs, when compared to those regulated by Nrf1α or Nrf1γ. However, Nrf1α contributed to a maximum ratio of the up-regulated to total DEGs, whereas Nrf1γ was rather attributable to a maximum ratio of the down-regulated to total DEGs.

**Figure 3.**
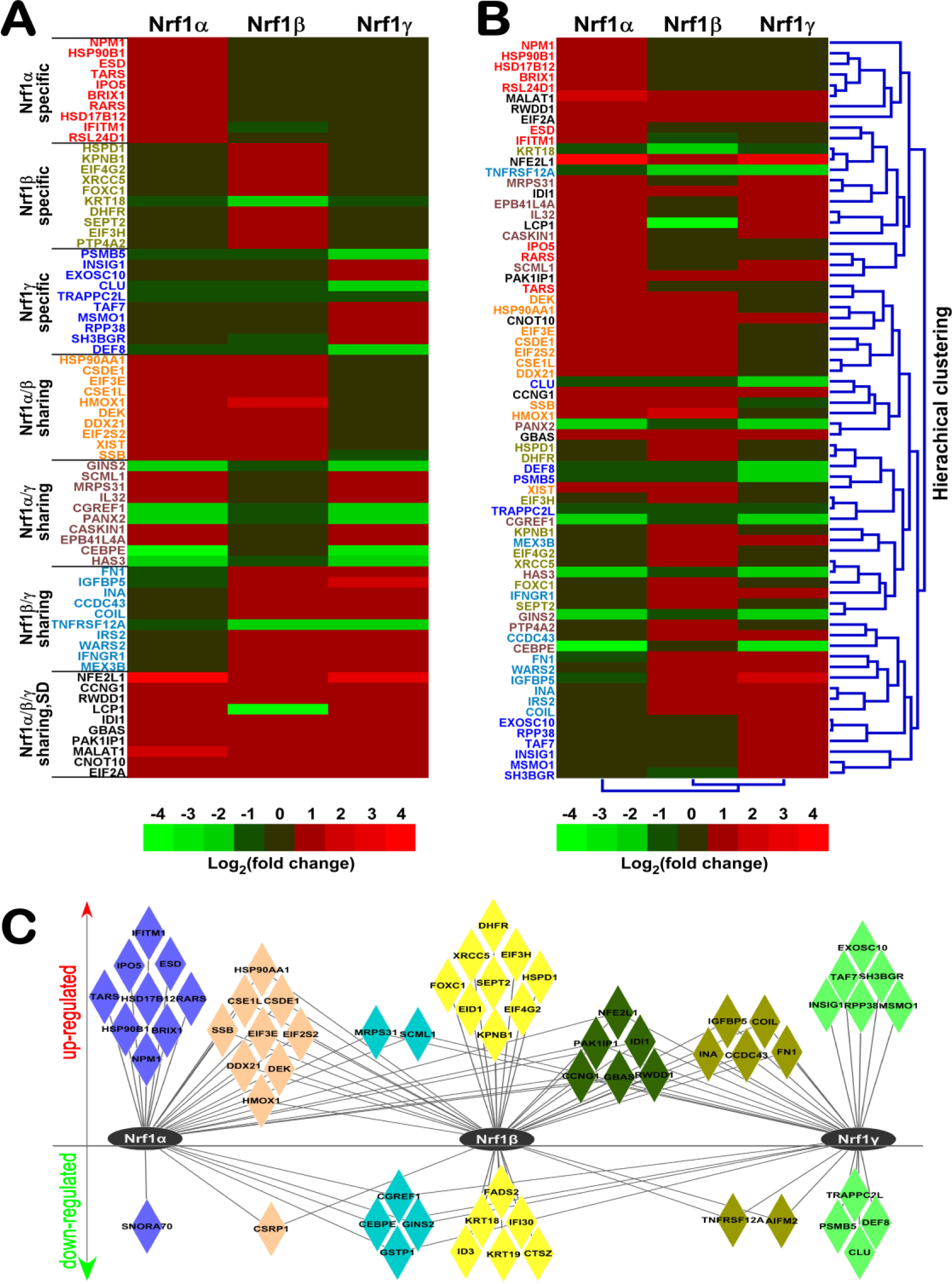
Expression patterns of top-ranked expression DEGs under regulated by three Nrf1 isoforms. (A, B) The heatmaps with distinct clusters were created for specific and/or sharing top 10 DEGs regulated by different isoforms of Nrf1. Differences in expression of distinct genes are shown in different colour backgrounds as scaled according to the Log_2_ (fold changes). SD (single difference), *LCP1.* (C) The network of the unique and/or common top-ranked DEGs amongst three sample groups after being normalized by control values.

### Interaction networks of Nrf1-related genes upon expression of its different isoforms

Since typical functions of genes are exerted in critical biological process through interaction networks, study of gene family interaction networks are useful to investigate potential gene functions [21]. As illustrated in Figure 4A, two Nrf1-mediated networks were constructed, in which some interactive proteins were identified by BioGRID (*upper three panels*) [22] and STRING (*lower three panels*) [23], respectively. Distinct expression profiles of these putative genes involved in the networks were extracted from distinct RNA-Seq data sets, which were reflected with different gradient colors in accordance with fold changes (Fig. 4A, and Tables S13 and S14). In the network identified with the BioGRID database, 12 of 19 genes showed a similar regulation trend, whereas other 7 genes displayed different regulation trends among three Nrf1 isoforms. Within these different regulation trends, three genes *MAFG, BRD9* and *FBXW7* were differentially regulated by Nrf1γ from other two isoforms, another three genes *RUSC2, CAPN1* and *HCFC1* were differentially regulated by Nrf1β, the remaining one gene *C8ORF33* was differentially regulated by Nrf1α. By comparison of the network identified with the STRING database, 5 of 11 genes showed an uniform regulation trend, whilst the other 6 genes were differently regulated: i) three genes *MAFG, NRF1* and *FBXW7* were differentially regulated by Nrf1γ from other two isoforms; ii) two genes *C3ORF35* and *MAFK* were differentially regulated by Nrf1β; and iii) only one gene *SP7* was differentially regulated by Nrf1α. Collectively, the interaction network analysis indicates that three isoforms of Nrf1 could diversely regulate its target genes. Such changes in the expression abundance of one isoform may also influence its overall transcriptional regulation of Nrf1-target genes. In fact, transcriptional expression of different subsets of target genes (driven by AP1-like AREs) was principally attributable to precision regulation by distinct functional heterodimers of CNC-bZIP family members (i.e. Nrf1, Nrf2, Nrf3, Bach1 and Bach2) with small Maf or other bZIP factors [24]. Such transcription of these factors was also further compared to determine distinct Nrf1-specific effects (Fig. 4B, and Table S15).

**Figure 4.**
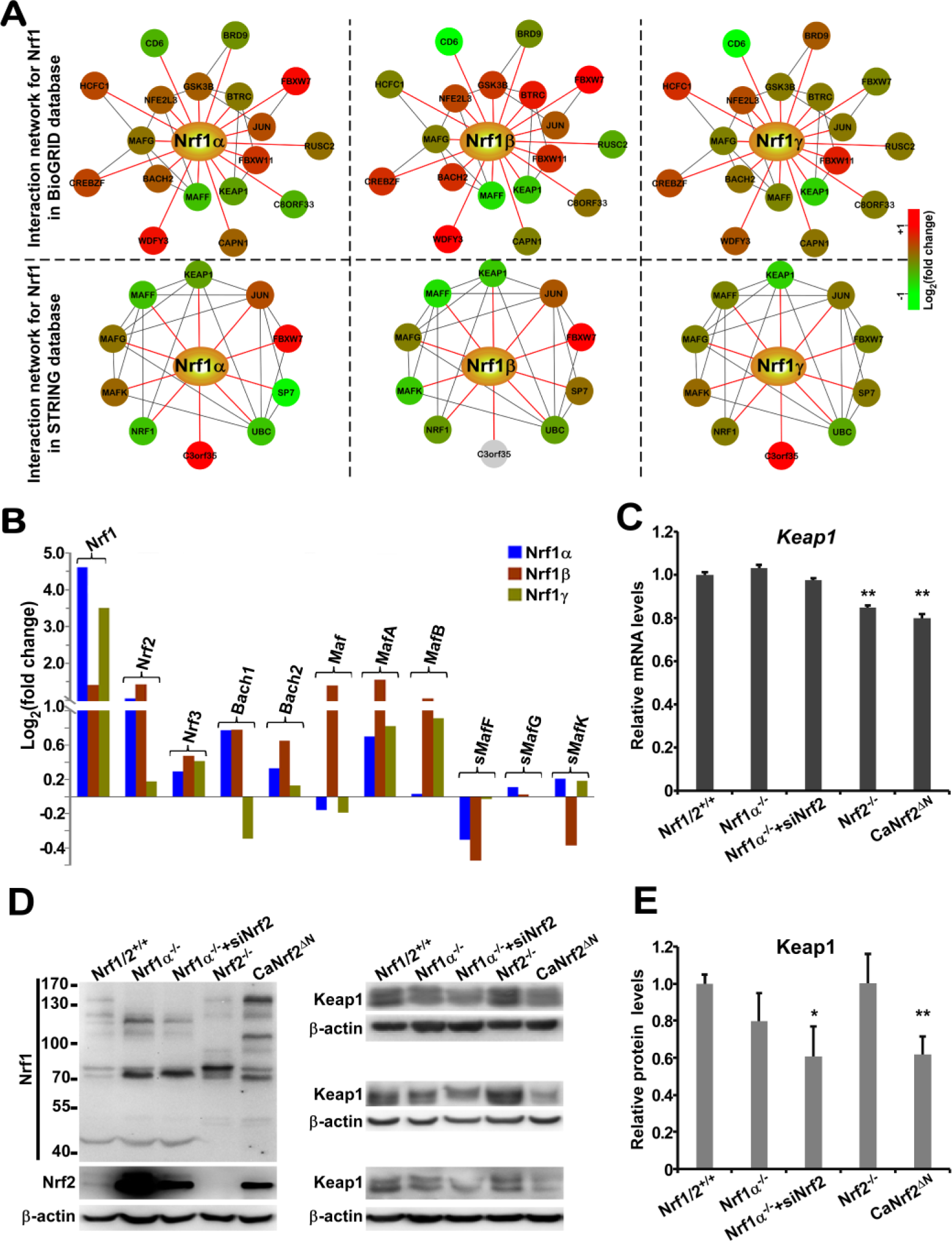
Nrf1-related genes in interaction networks and their expression regulated by different Nrf1 isoforms. (A) Two interaction networks for Nrf1 were established on the base of the BioGRID database and STRING database. The interactor nodes showing up-regulation or down-regulation were marked in red or green, respectively. Such genes were marked with gradient colours from green to red according to the log2 fold changes, but gray indicates that a gene was not expressed. (B) The expression comparison of Nrf1, Nrf2 and Nrf3, Bach1 and Bach2, together with small Maf families. (C to E) Expression of Keap1 (Kelch-like ECH-associated protein 1) at its mRNA levels (C) and protein levels (D, E), was determined in the presence or absence of Nrf1α and/or Nrf2. The error bars indicate mean ± SD (n = 3), with significant decreases (*p < 0.05, **p < 0.01) being calculated relative to the corresponding control values.

Intriguingly, the N-terminal Neh2 (Nrf2-ECH homology 2) domain of Nrf2, which contains a redox-sensitive Keap1 (Kelch-like ECH-associated protein 1)-binding degron targeting the homoeostatic CNC-bZIP protein to ubiquitin ligase cullin-3/Rbx1-dependent proteasomal degradation pathway [25–28], is represented by the Neh2-like region within Nrf1α [24], but the latter CNC-bZIP protein is not regulated by Keap1 [10]. This notion also appeared to be further supported by the observation that both mRNA levels (Fig. 4C) and protein levels (Fig. 4, D & E) of Keap1 were differentially expressed under distinct genetic backgrounds of knockout (i.e. *Nrf1α*^*-/-*^ or *Nrf*^*-/-*^), siRNA (i.e. si N rf2) or constructive activation of Nrf2 mutant (i.e. caNrf2^ΔN^). In turn, the results also reveal that significant changes in the expression of Keap1 are accompanied by altered expression of Nrf2, but not Nrf1α. This implies a feedback circuit existing between Nrf2 (rather than Nrf1) and its negative regulator Keap1.

### Distinct effects of Nrf1 isoforms on proteasomal expression in response to stimulation by proteasomes inhibitor

The ubiquitin-proteasome system (UPS) is crucial for eukaryotic cells to adjust its capacity of protein degradation to changing proteolytic requirements, because these proteins are marked with polyubiquitin chains targeting for their degradation by the 26S proteasome, an ATP-dependent complex consisting of the 20S proteolytic particle capped by one or two of the 19S regulatory particles [29]. Importantly, coordinated regulation of proteasomes by Nrf1, but not Nrf2, occurs through proteasome-limited proteolytic processing of the former CNC-bZIP protein into a mature active factor to mediate Nrf1-target proteasomal gene expression in the ‘bounce-back’ response to relative lower doses of proteasomal inhibitors [30–33]. Nrf1-mediated induction of proteasomes subunits results in significant increases in mRNA expression levels of all proteasomal subunits only upon exposure to lower concentrations of proteasomal inhibitors, but this feedback compensatory response is prevented by high concentrations of proteasomal inhibitors [33]. Therefore, there exists a bidirectional regulatory feedback circuit between Nrf1 and the proteasome [24, 31–34]. However, our data revealed that almost none of proteasomal subunits were differentially expressed at basal levels, because they were unaffected by stably-induced expression of any one of Nrf1 isoforms (Fig. 5A, and Table S16).

**Figure 5.**
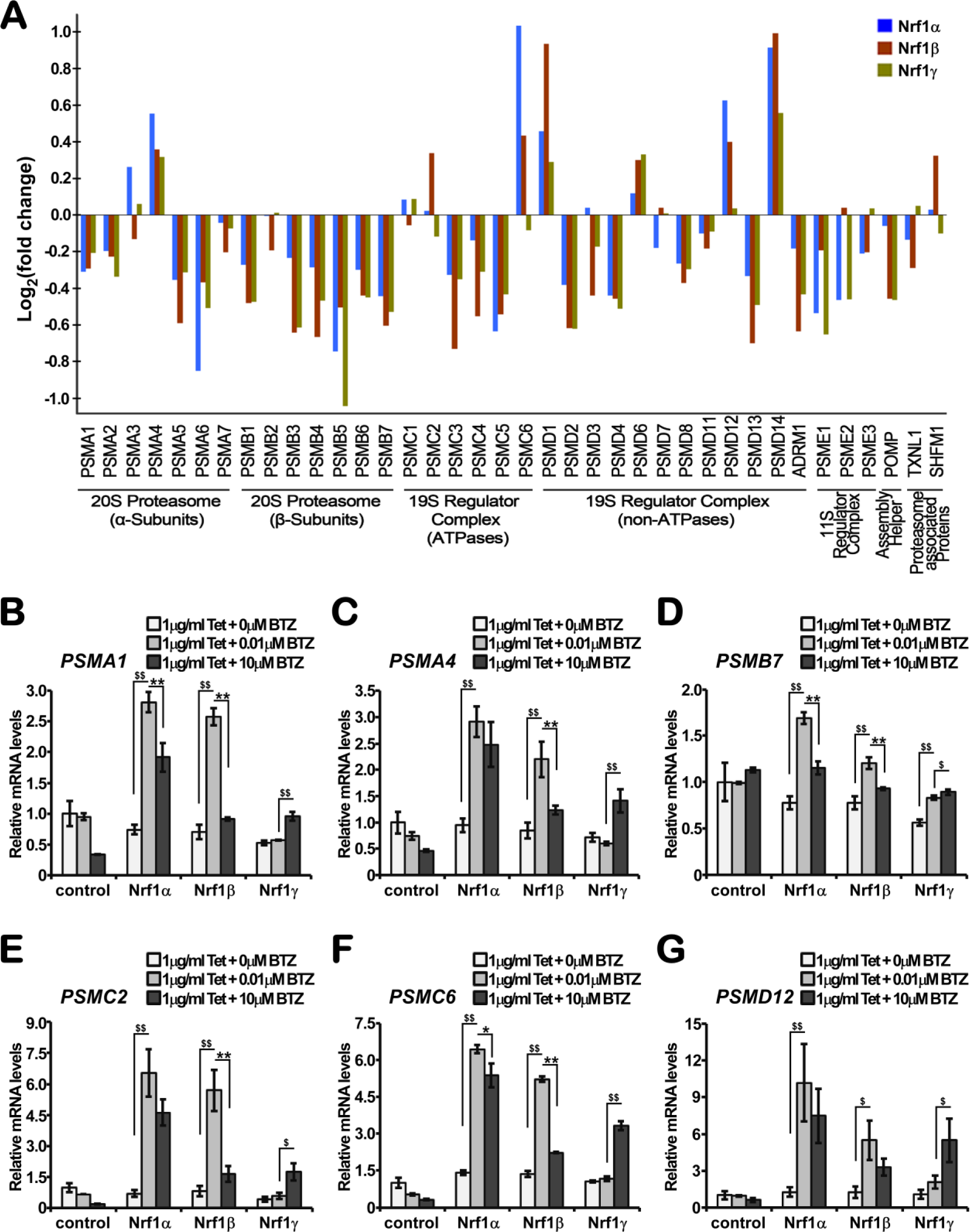
Distinct effects of Nrf1 isoforms on proteasomal expression in distinct responses to its inhibitor. (A) The RNA-Seq levels of proteasomal subunit genes were selected for distinct isoform-specific regulation relative to control values. (B to G) When compared with experimental controls, Nrf1α-, Nrf1β-and Nrf1γ-expressing cells were treated with 1 μg/ml of Tet alone or plus a lower (0.01 μmol/L) or higer (10 μmol/L) doses of bortezomib (BTZ) for l6 h. The regulatory effects of Nrf1 isoforms, along with distinct concentrations of BTZ, on 26S proteasomal gene transcription were then analyzed by real-time qPCR. The data are represented by a mean ± SD of three independent experiments, with significant increases ($p < 0.05, $$p < 0.01) and significant decreases (*p < 0.05, **p < 0.01) being determined, relative to their corresponding control values.

To address this, we validated Nrf1-stimulated induction of proteasomal subunits by its inhibitors. Experimental cells that had treated with 1 μg/ml of Tet alone or plus a low (0.01 μmol/L) or a high (10 μmol/L) concentration of bortezomib (BTZ) for 16 h were subjected to further determination of mRNA expression levels of some proteasomal subunits, including *PSMA1, PSMA4, PSMB7, PSMC2* and *PSMD12* by real-time qPCR (Fig. 5, B to G). As expected, the results demonstrate that all the mRNA levels of these proteasomes examined were increased following exposure of Nrf1α-or Nrf1β-expressing cells to 0.01 μmol/L of BTZ. Conversely, such increased expression was significantly prevented by additional exposure of Nrf1α-or Nrf1β-expressing cells to 10 μmol/L of BTZ. Intriguingly, the parallel experimentations of Nrf1γ-expressing cells revealed that almost no increases in the BTZ-stimulated expression of those aforementioned proteasomes except *PSMB7* were detected, even upon exposure to the low concentration at 0.01 μmol/L of BTZ, as consistent with the notion that Nrf1γ is likely to act as a dominant-negative inhibitor of Nrf1. However, it is full of curiosity about the finding that the high concentration (10 μmol/L) of BTZ enabled for significant stimulation of Nrf1γ to increase mRNA expression levels of all examined proteasomal subunits, albeit the detailed mechanism(s) remains to be further explored

### Different regulatory effects of distinct Nrf1 isoforms on the downstream target genes

To confirm the conclusion drawn from the RNA-Seq data, some downstream genes of Nrf1 were selected for further experimental validation by qRT-PCR analysis (Fig. 6). Such known Nrf1-target genes included those encoding HO1 (heme oxygenase 1) [35–37], GCLM (glutamate-cysteine ligase, modifier subunit), GCLC (glutamate-cysteine ligase, catalytic subunit) [38–41], MT1E (metallothionein 1E) [42], PGC-1β (peroxisome proliferator-activated receptor gamma, coactivator 1 beta) [43] and LPIN1 (lipin1) [44]. As expected, the experimental results revealed that these Nrf1-target genes exhibited different trends to be expressed specifically in distinct isoform-expressing stable cells (Fig. 6, D to H), each of which is distinctive from those measured in the other two isoforms-expressing cell lines, but with an exception that *HO1* was up-regulated by all three isoforms (Fig. 6C).

**Figure 6.**
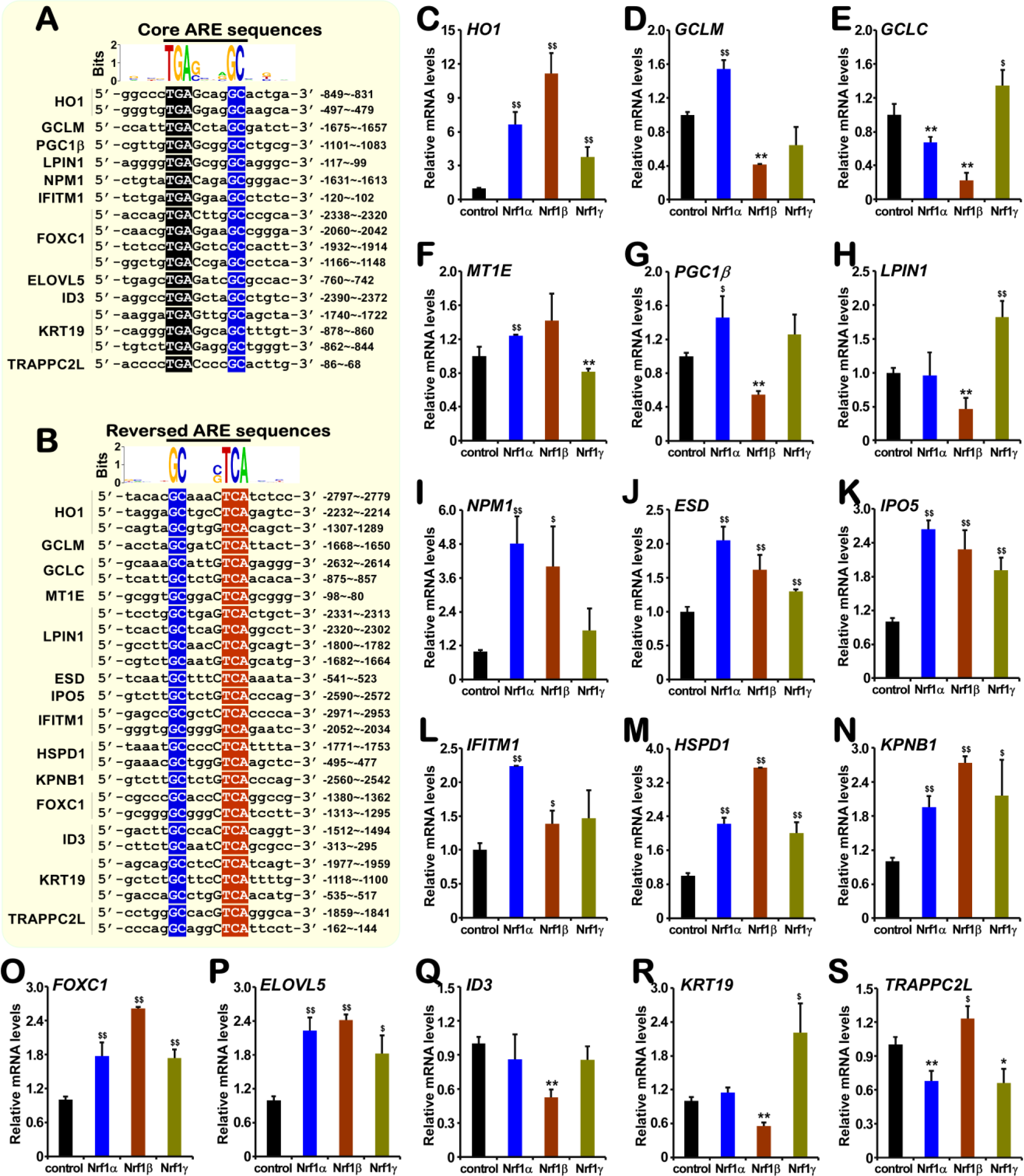
Distinct isoform-specific regulation of Nrf1-target ARE-driven genes. (A, B) The core and reversed ARE sequences are encompassed in the promotor regions of these Nrf1-regulated genes. When compared with experimental controls, Nrf1α-, Nrf1β-and Nrf1γ-expressing cells were induced with 1 μg/ml of Tet for 12 h. (C to H) Expression of those known Nrf1-target downstream genes, such as *HO1, GCLC, GCLM, MT1E, PGC1P* and *LPIN1*, was determined by real-time qPCR. (I to S) Expression of other potential downstream genes of Nrf1 that were selected on the base of RNA-Seq data was further validated by real-time qPCR. The data are represented by a mean ± SD of three independent experiments, with significant increases ($p < 0.05, $$p < 0.01) and significant decreases (*p < 0.05, **p < 0.01) being indicated, relative to their corresponding controls.

Next, other ARE-driven downstream genes were also selected, to be potentially regulated by Nrf1, based on the RNA-Seq data according to RPKM and log2 fold change, and then determined by real-time qPCR. As shown in Figure 6 (I to L), those genes encoding NPM1 (nucleophosmin 1), ESD (esterase D), IPO5 (importin 5) and IFITM1 (interferon induced transmembrane protein 1) were differentially up-regulated by Nrf1α. They were also significantly increased by Nrf1β, albeit their increased levels appeared to be less than those regulated by Nrf1α. It is, therefore, inferred that Nrf1β is not a dominant-negative inhibitor competitively against Nrf1α, this is consistent with the notion previously reported by our group [12, 45–47]. Furthermore, it is, to our surprise, that mRNA levels of both *ESD* and *IPO5* were significantly increased by Nrf1γ (Fig. 6, J & K), even though their fold changes were less than those measured from the other two cases of Nrf1α and Nrf1β.

The additional four genes, such as *HSPD1* (heat shock 60kDa protein 1), *KPNB1* (karyopherin subunit beta l), *FOXC1* (forkhead box C1) and *ELOVL5* (ELOVL fatty acid elongase 5) were differentially up-regulated by Nrf1β, at less increased levels than equivalents regulated by Nrf1α and Nrf1γ (Fig. 6, M to P). Moreover, both *ID3* (inhibitor of DNA binding 3, dominant negative helix-loop-helix) and *KRT19* (keratin 19) were differentially down-regulated by Nrf1β (Fig. 6, Q & R), whilst *KRT19* was up-regulated by Nrf1γ. Lastly, the gene *TRAPPC2L* (trafficking protein particle complex 2-like) was down-regulated by Nrf1α and Nrf1γ, but up-regulated by Nrf1β (Fig. 6S). Notably, these genes are all known to exert their respective biological functions and be involved in different pathways (see Tables S5 to S10). Further bioinformatic search demonstrated that all the above-described putative Nrf1-target genes contains more than one of the highly-conserved ARE motifs and/or its reversed sequences (Fig. 6, A & B). Thus, it is inferred that these ARE-driven genes are likely to be differentially regulated by distinct Nrf1 isoforms, but this warrants further experiments to elucidate which ARE motifs are functionally responsible for an Nrf1-specific isoform to regulate a given gene.

## Discussion

Differential expression of different subsets of cognate genes is dependent on their enhancers and promoter regions containing distinct *cis*-regulatory consensus sequences (e.g. AP-1-like AREs) [24]. Transcriptional expression of AREs-driven genes is thus determined by differential recruitment of Nrf1, Nrf2 and Nrf3, in different combinations with each of their heterodimeric partners (e.g. sMaf, c-Jun, JunD or c-Fos), to target gene promoters. Of note, Nrf1 and Nrf2 are two important CNC-bZIP transcription factors expressed ubiquitously in various vertebrate tissues and thus elicit their putative combinational or competitive functions. Relative to the well-known water-soluble Nrf2, less attention has hitherto been drawn to the membrane-bound Nrf1 [24]. However, major discoveries that had been made in the past twenty-five years have revealed that Nrf1, but not Nrf2, has been shown to be indispensable for maintaining cellular homoeostasis and organ integrity during normal development and healthy growth, as well as a vast variety of other patho-physiological processes. Importantly, several significant pathological phenotypes were developed in different transgenic mice (expressing distinct mutants of loss-of-function of Nrf1), including embryonic lethality, fetal anemia, lipid metabolic disorder, obesity, fatty liver, NASH, liver cancer, neurodegenerative diseases, hyperinsulinemia, diabetes, Warbug effect with high glycolysis. In addition, other mice expressing gain-of-function mutants of Nrf1 displayed glucose metabolic disorder, insulin resistance, diabetes and reduced body-weight. Thus it is inferable that the functional activity of Nrf1 is finely tuned, to a robust homeostatic extent, by a steady-state balance between its production and the concomitant processing into distinct isoforms before being turned over, which are together coordinated to confer on the host cytoprotection against a variety of cellular stresses.

Accumulating evidence has unraveled that over eleven of distinct Nrf1 isoforms are produced from the single *Nfe2l1/Nrf1* gene, though differentially expressed, in different mammalian species. These isoforms are synthesized by translation through distinct initiation signals (i.e. the first or internal start ATG codons) embedded in different lengths of open reading frames, portions of which can be alternatively spliced from intact or longer transcripts. The resulting variations in the abundance of each isoform may be not only influence the whole transcriptional functions of Nrf1 to regulate distinct subsets of cognate target genes and also contributes to the nuance in between distinct pathological phenotypes [24]. Therefore, it is of crucial important to determine differences in transcriptional regulation of cognate genes mediated by each Nrf1 isoform. Although this is hard, our present study has identified differential expression profiles of distinct target genes regulated by Nrf1α, Nrf1β and Nrf1γ alone or in their cooperation respectively. Here, we have also determined differences in the transcriptional regulation of Nrf1-target genes by between each Nrf1 isoforms. Notably, Nrf1α and Nrf1β are two major isoforms contributing to the main Nrf1-mediated transcription of downstream genes at RNA levels, such that a vast majority of differentially expressed genes are up-regulated by Tet-inducible expression of these two isoforms. On the contrary, stably Tet-inducible expression of Nrf1γ as a putative dominant-negative inhibitor is likely to interfere with the functional assembly of active transcription factors (Nrf1α, Nrf1β, and even Nrf2), leading to down-regulation of several key genes, some of which are up-regulated by Nrf1α and Nrf1β. These findings are consistent with our previous reports [12, 45–47]. Collectively, these findings are very helpful to elucidate which isoforms of Nrf1 contribute to different transcription of distinct subsets of target genes that are involved in those significant pathological phenotypes. Thus, this study has provided three cell models to facilitate the future development of Nrf1 isoform-specific targets for chemoprevention against relevant diseases (i.e. cancer and diabetes).

Furthermore, it should also be noted that we have presented most of the data about Nrf1γ in the present study to further support our previous notion [12, 45–47], which revealed this low molecular weight isoform acts as a dominant-negative inhibitor of Nrf1 competitively against the functional heterodimeric assembly of either an active transcription factor (i.e. Nrf1α, Nrf1β, even Nrf2 and Nrf3) or another homologous trans-repressor (i.e. Bach1 and Bach2) with one of their cognate partner sMaf or other bZIP proteins (e.g. c-Jun, c-Fos, or ATF4). Therefore, it is plausible that either transactivation or transrepression of distinct subsets of similar and/or different target genes driven by AP1-like ARE/EpRE batteries is dependent on a nuance between different temperospatially-assembled heterodimeric complexes of these CNC-bZIP factors with their partners. This is to say, understandably, that the dominant-negative form of Nrf1γ is much likely to counteract (or interfere with) the putative activity of its prototypic factors Nrf1α/β to transactivate or transrepress their downstream genes. For this reason, it is thus deduced that if some genes are down-regulated by Nrf1α/β, this down-regulation is abolished or even reversed to allow for their activation by Nrf1γ. In addition, there also exists an exception that a few number of Nrf1-target genes (as shown in Fig. 6) are positively regulated by Nrf1γ, seemingly as done by Nrf1α/β. It cannot be theoretically ruled out that Nrf1γ might also have a similar potential effect to that elicited by sMaf factors, which can still form a functional heterodemeric assembly with an activator albeit it lacks a *bona fide* transctivation domain, but this remains to be further determined in the future experiments.

## Materials and Methods

### Chemicals and antibodies

All chemicals were of the highest quality commercially available. Hygromycin-B and blasticidin were purchased from Invitrogen Ltd, which served as double screening drugs to select the positive clones by final concentrations of 150 μg/ml and 15 μg/ml, respectively. The inducible reagent tetracycline hydrochloride was from Sangon Biotech Co (Shanghai, China) and used at a final concentration of 1 μg/ml. The proteasome inhibitor bortezomib was purchased from ApwxBio (USA). The antibody against endogenous Nrf1 proteins was acquired from our lab (i.e. Zhang’s indicated in this study [48]). Moreover, mouse monoclonal antibody against the V5 epitope was from Invitrogen Ltd, whilst β-actin and secondary antibodies were obtained from Zhongshan Jinqiao Co (Beijing, China).

### Expression constructs and cell culture

The cDNA fragments encoding three Nrf1 isoforms Nrf1α, Nrf1β and Nrf1γ were cloned into pcDNA5/FRT/TO-V5 expression vector, before being transfected into the host cell line Flp-InTMT-RExTM-293. The empty expression vector-transfected host cell served as a negative control. The positive isoform-expressing clones were selected by using 150 μg/ml hygromycin B and 15 μg/ml blasticidin and then stable expression of Nrf1α, Nrf1β and Nrf1γ were induced by addition of tetracycline. All cell lines used in this study were cultured in DMEM medium supplemented with 10% FBS and maintained in the 37°C incubator with 5% CO_2_.

### Western blotting

Experimental cells were harvested in a lysis buffer (2 mM Tris pH 7.5, 5 mM NaCl, 0.5 mM Na_2_EDTA, 0.04 mM DTT, 0.5% SDS) containing 2 μg/ml protease inhibitor cocktail (Roche, Germany). The protein concentration of lysates was quantified by using BCA Protein Assay Reagent (Bi-Yintian, Beijing, China). Equal amounts of protein prepared from cell lysates were loaded into each electrophoretic well so as to be separated by SDS-PAGE, followed by visualization by immunoblotting with V5 antibody as described previously [49], and β-Actin was served as an internal control to verify amounts of proteins that were loaded in each well.

### The RT-qPCR Analysis

Total RNAs were extracted from experimental cells by using an RNAsimple total RNA kit (Tiangen, Beijing, China). Then, 1.5 μg of total RNAs were used as a template for the subsequent synthesis of cDNA by using a RevertAid first strand cDNA synthesis kit (Thermo Fisher Scientific, USA). The resulting cDNA products (15 ng) served as the templates of quantitative real-time PCR within 5 μl of the MixGoTaq®qPCR Master Mix (Promega, USA). Each of RT-qPCR with distinct pairs of primers (listed in the supplementary Table S17) was performed in the following conditions: activation at 95°C for 30s, followed by 40 cycles of 10s at 95°C, and 30s at 60°C. These PCR reactions were carried out in at least 3 independent experiments that were each performed triplicate. The comparative Ct method was employed for quantification of indicated mRNA expression levels before being normalized to β-Actin.

### Bioinformatics analysis of RNA-Seq data

Each isoform-expressing cell lines (i.e. Nrf1α, Nrf1β and Nrf1γ), along with the negative control cells, were allowed for growth in 6-well plates, before being induced by tetracycline (1 μg/ml) for 12 h. Total RNAs were extracted by using an RNAsimple total RNA kit (Tiangen, Beijing, China) and the integrity of RNAs was checked by an Agilent Bioanalyzer 2100 system (Agilent technologies, Santa Clara, CA). Subsequently, RNA-Seq was carried out by Beijing Genomics Institute (BGI, Shenzhen, China) on an Illumina HiSeq 2000 sequencing system (Illumina, San Diego, CA) after the sample library products are ready for sequencing.

After examining the sequence quality and removing the “dirty” raw reads, which contain low quality reads and/or adaptor sequences, clean reads (the high quality sequences after data cleaning) were generated and stored as FASTQ format [50]. Then, clean reads were mapped to the reference human genome by using SOAP2 [51], and gene expression levels were calculated by using the RPKM (Reads Per Kilobase of feature per Million mapped reads) method [16]. The gene expression regulated by each Nrf1 isoforms, relative to the control sample, was calculated as Log_2_ (fold change), with a *P*-value corresponding to differential gene expression test and FDR (False Discovery Rate), which is a method to determine the threshold of P-value in multiple tests [17, 52]. Both FDR ≤ 0.001 and the absolute value of Log_2_ (fold change) ≥ 1 were taken as the threshold to identify differentially expressed genes. To give a better understanding of potential functions of the DEGs, both GO and KEGG pathway analysis were performed by using the online tools DAVID (https://david.ncifcrf.gov/) and KEGG (http://www.kegg.jp/) databases, respectively. In addition, the putative interaction networks of Nrf1-related genes were searched from the databases of BioGRID (https://thebiogrid.org/) and STRING (https://string-db.org/), before being annotated with sequencing data by the Cytoscape software [53].

### Statistical analysis

The data provided in this study were represented as the mean ± SD and were analyzed using the Student’s *t*-test or Fisher’s exact test, as appropriate. The value of *p* < 0.05 was considered as a significant difference.

## Competing interests

The authors declare no competing financial interests.

## Author contributions

M.W. performed most of bioinformatic analyses and experiments, collected the resulting data and prepared draft of this manuscript with most figures and supplementary tables. L.Q. helped M.W. together with molecular cloning to create expression constructs and performed western blotting of Keap1. X-F.R. prepared the RNA-sequencing samples. Y-J.S. helped M.W. with functional annotation of differentially expressed genes. Y-G.Z. designed this study, analyzed all the data, helped to prepare all figures, wrote and revised the paper.

## Acknowledgments

The study was supported by the National Natural Science Foundation of China (key programs 91129703, 91429305 and project 31270879) awarded to Prof. Yiguo Zhang (University of Chongqing, China), and in part funded by

Chongqing University postgraduates’ innovation project (No. CYB15024) awarded to Mr. Lu Qiu.

